# Identification of Altered Potassium Channels for Drug Repurposing in Long COVID Patients

**DOI:** 10.64898/2026.06.18.733062

**Authors:** John P. George, Kiran Bharat Gaikwad, Jyoti Sharma

## Abstract

Long COVID (LC) is a complex condition characterized by persistent, chronic multisystem manifestations, with a significant proportion of patients exhibiting neurological symptoms. Human ion channels (HICs), particularly potassium channels, are abundantly expressed in the nervous system and linked to key metabolic processes, making them potential candidates for understanding LC pathophysiology and drug repurposing. Meta-analysis of RNA-Seq datasets from COVID-19-recovered and LC patients was performed to identify altered HICs in LC. Differential gene expression analysis, functional enrichment analysis, and weighted gene co-expression network analysis were performed to uncover key genes, pathways, and co-expression modules consisting of HICs, lipid metabolism-, and immune signaling-related genes. A total of 715 dysregulated genes, including eighteen HICs were identified, among which seven were potassium channels. Three significant modules containing HICs, lipid metabolism-, and immune signaling-related genes were identified and found to be associated with antigen processing and presentation, complement and coagulation cascades, and cytokine-related pathways. Additionally, drug-gene interaction analysis led to identification of approved drugs targeting *KCNA6*, *KCNJ10*, *KCNN3*, and *KCNH4* that might provide opportunities for drug repurposing in neurological manifestations in LC. Further experimental validation is required to establish their efficacy and assess their potential for translation into clinical applications for patients with LC.

## 1 Introduction

Six years after the COVID-19 pandemic, long COVID (LC) remains a significant concern among individuals with a history of SARS-CoV-2 infection. It affects a substantial proportion of COVID-19 survivors worldwide, presents with overlapping symptom subtypes and is more likely to occur in individuals with severe disease, comorbidities, and certain demographic risk factors (1). It is primarily characterized by persistent fatigue, brain fog, dyspnea, chest pain, palpitations, and other multisystem manifestations involving the neurological, respiratory, cardiovascular, and musculoskeletal systems (1), (2). These symptoms are reported to arise from multiple overlapping mechanisms, including viral persistence, immune dysregulation, chronic inflammation, microvascular dysfunction, autonomic imbalance, and reactivation of latent viruses (2).

Significant alterations in host gene expression are induced during COVID-19 infection, primarily dysregulating immune response genes, proinflammatory cytokines, and interferon (IFN) pathways (3). Emerging studies using transcriptomic profiling have revealed persistent alterations in gene expression associated with immune dysregulation, inflammation, and metabolic imbalance in affected individuals with elevated proinflammatory cytokines, including interleukin-6 and tumor necrosis factor-α, and altered T and B-cell profiles (4). Elevated expression levels of IFNs in patients with severe COVID-19 have been reported to decline over time in LC patients, suggesting cytokine exhaustion as a possible driver of LC (5). More integrated transcriptomic analysis is required to elucidate the interconnected molecular networks driving LC pathophysiology that could also provide therapeutic strategies in LC management.

Human ion channels (HICs) play a crucial role in regulating cellular homeostasis, membrane potential, and signal transduction and have been increasingly reported in the pathophysiology of viral infections, including COVID-19 (6). Several studies have highlighted the dysregulation of HIC activity during SARS-CoV-2 infection, suggesting their involvement in COVID-19-associated complications (7), (8). They contribute to critical processes including viral entry, replication, and host immune responses, thereby influencing disease progression and severity (9).

Of these, potassium channels constitute a major class of HICs that are essential in maintaining ionic balance, regulating membrane repolarization, and controlling cellular excitability across neuronal, cardiac, and epithelial systems (10), (11), (12), (13). Dysregulation of potassium channels is known to disrupt K⁺ homeostasis and membrane potential, leading to altered neuronal signaling and impaired cellular function (14), (15), (13). Hypokalemia, characterized by reduced serum potassium levels, is a serious condition that can lead to electrolyte imbalance, arrhythmias and seizures and has been frequently reported in patients with acute COVID-19 (16), (17). Notably, emerging case reports have documented persistent hypokalemia beyond the acute phase of infection, indicating sustained disruption of potassium homeostasis and a potential link to LC (17), (18). These channelopathies are associated with neuropsychiatric manifestations, cardiac abnormalities, and immune dysregulation and may also facilitate viral entry and persistence, highlighting their potential role in the multisystem nature of LC (19), (20), (21). Moreover, due to their accessibility and functional significance, HICs have emerged as promising druggable targets (9). However, their specific role and regulatory dynamics in the context of LC remain largely unexplored.

Additionally, HICs, lipid metabolism and immune signaling pathways play critical roles in the host response to viral infections and the development of persistent disease states. Lipid metabolism is essential for viral replication, membrane remodeling, and energy homeostasis, while dysregulation of immune signaling pathways contributes to the chronic inflammation and prolonged symptomatology observed in LC (22), (23), (24). Emerging evidence suggests that these biological processes do not act independently but are highly interconnected, with significant cross-talk influencing disease progression (22), (23). However, the integrated interactions between HICs, lipid metabolism, and immune signaling pathways have not been systematically explored in LC. Understanding these interconnected molecular networks may provide deeper insights into the mechanisms underlying disease persistence.

Given the interconnected nature of these molecular processes, a systems-level approach is essential to understand LC pathophysiology. Traditional differential expression analyses provide valuable insights into individual genes but often fail to capture the coordinated interactions between them. In contrast, gene co-expression network analysis enables the identification of functionally related gene modules and underlying regulatory patterns, offering a more holistic view of disease-associated molecular mechanisms (25). Such approaches may reveal key genes, thereby facilitating the identification of critical molecular drivers in LC.

With the lack of approved targeted therapies for LC, there is a need to identify effective treatment strategies. Drug repurposing offers a time- and cost-efficient approach by utilizing existing approved drugs with known safety profiles (26). Integrating transcriptomic findings with drug–target interaction data can facilitate the identification of potential therapeutic candidates by linking disease-associated molecular alterations to actionable targets (26), (27). This may accelerate the translation of molecular insights into clinical applications, providing a practical framework for the management of LC.

## 2 Methodology

### 2.1 Data Acquisition and Processing

RNA sequencing (RNA-Seq) data from 62 COVID-19-recovered and 70 LC patients were obtained for this study. Two datasets (GSE251849 and GSE226260) were downloaded from the Gene Expression Omnibus database, whereas one dataset (PRJNA1184005) was acquired from NCBI’s BioProject. A list of 493 HICs was downloaded from the HUGO Gene Nomenclature Committee database. Lipid metabolism-related genes (LMGs) were collected from the literature (28), (29), (30), (31), (32), (33), (34), (35), (36), (37), (38). Similarly, a list of immune signaling-related genes (ISGs) was obtained from the InnateDB (39). The quality of the RNA-Seq data was assessed using FastQC (v. 0.12.1) and further processed using Trimmomatic (v. 0.39) (40). Quality filtered reads were aligned to the human genome GRCh38.p13 using STAR aligner (v. 2.7.11b) followed by gene expression quantification using HTSeq (v. 2.1.2) (41), (42). Batch effects arising from technical variability across samples were corrected using the ComBat_seq function of the sva package (v. 3.54.0) prior to downstream analyses (43). The overall workflow is summarized in Figure 1.

**FIGURE 1.**
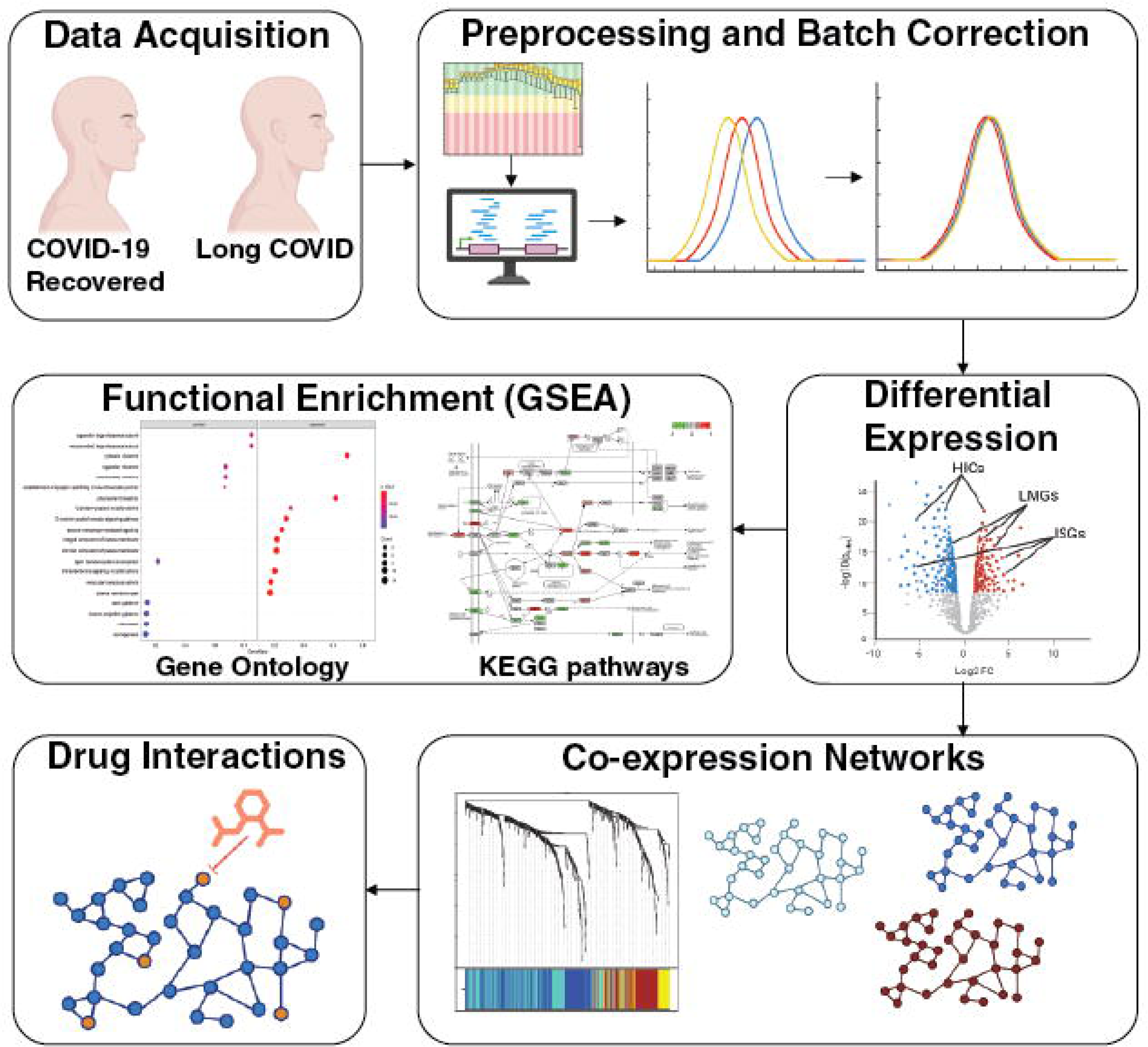
A schematic overview of the several analyses carried out to identify potential altered potassium channels. Publicly available transcriptomic data of COVID-19-recovered and long COVID was downloaded, processed, and treated for batch effect removal. Differential expression analysis was performed to identify differentially expressed genes. Functional enrichment for the identified differentially expressed genes (DEGs) was performed to identify associated Gene Ontology terms and pathways. Gene co-expression network analysis was performed to identify significant clusters comprising differentially expressed human ion channels (HICs) and lipid metabolism- and immune signaling-related genes. Approved drugs interacting with these critical HICs were identified. These drug-HICs interactions may provide potential strategies for drug repurposing.

### 2.2 Differential Expression

Differential gene expression analysis between COVID-19-recovered and LC gene expression profiles was performed using the DESeq2 package (v. 1.46.0) of R (R Foundation for Statistical Computing, Vienna, Austria) (44). Genes with a *p-*value <= 0.05 and |log2fold change| > 0.6 were classified as differentially expressed genes (DEGs). A volcano plot was generated using the ggplot2 package (v. 3.5.2), with differentially expressed HICs, LMGs, and ISGs specifically labeled.

### 2.3 Functional enrichment analysis

The matrix generated from the differential gene expression analysis was used to perform functional enrichment analysis using ClusterProfiler (v4.14.6), an R package (45). Significant (*p*-value ≤ 0.05) enriched Gene Ontology (GO) terms and KEGG pathways that were activated (up-regulated) or suppressed (down-regulated) in LC patients were identified using the gseGO and gseKEGG functions, respectively. Further, the processes and pathways were visualized using the dotplot function implemented in R.

### 2.3 Network Construction and Pathway Analysis

Co-expression networks were constructed using the weighted gene co-expression network analysis (WGCNA) package (v. 1.74) of R (46). Expression profiles of DEGs were provided as an input and transformed using DESeq2’s variance stabilizing transformation. To generate a scale-free topology network, a soft-thresholding power of 4 was selected. A minimum module size of 30 genes was used for module detection. After converting expression data into a topological overlap matrix (TOM), the dynamic tree-cutting approach was used to detect initial modules. Modules were merged based on substantially similar expression profiles of the genes. Eigengene-trait correlation analysis was performed to identify modules significantly associated with COVID-19-recovered and LC, and modules with a p-value ≤ 0.05 were considered significant. Gene significance and module membership (MM) values were utilized to assess the correlation between gene expression profiles, module eigengenes, and binary characteristics. Gene significance versus MM plots was used to assess intramodular gene relationships (46). The most significantly correlated modules were chosen, and the resulting networks were exported to Cytoscape (v. 3.10.3) using a weight threshold of 0.02 (47). Furthermore, pathways associated with the significant modules were identified using EnrichR (48). The R script used for the WGCNA analysis has been made publicly available in the GitHub repository, and the corresponding link is provided in the Data Availability section.

### 2.4 Drug Interactions

HICs in the co-expression networks were considered as potential druggable targets for drug repurposing. Approved drugs interacting with these HICs were identified using the Drug-Gene Interaction Database (DGIdb) (49). DGIdb provides curated drug–gene interaction records to allow hypothesis generation and the identification of potential therapeutic candidates for drug repurposing (49). To complement this target-based approach, transcriptome-based drug repurposing analysis was performed using the Connectivity Map (cMAP) CLUE platform (50). The differential gene expression signature comprising both up-regulated and down-regulated genes identified in LC patients was used as a query against the Touchstone reference database, employing the L1000 platform for high-throughput gene expression profiling. Connectivity between the LC gene expression signature and perturbagen-induced transcriptional profiles was computed using a weighted Kolmogorov-Smirnov enrichment statistic to generate connectivity scores for each disease-drug pair (51). A negative normalized connectivity score indicates that the perturbagen denoted an inverse transcriptional signature relative to the disease-associated gene expression profile, suggesting its potential to reverse disease-associated molecular alterations and serve as a candidate for drug repurposing (51), (52). Perturbagens with an fdr_q_nlog10 value greater than 1.3 (equivalent to FDR < 0.05) were considered statistically significant.

### 2.5 Validation of Identified DEGs in Independent Datasets

RNA-Seq data from 35 post-COVID-19 individuals from GSE222253 along with a longitudinal RNA-Seq data from 50 blood samples (25 samples each from the 3-6 month and 12–15-month post-COVID-19 infection time points) from GSE267625 were utilized to validate the identified differentially expressed HICs (53).

## 3 Results

### 3.1 Differential Gene Expression and Functional Enrichment

The total 715 DEGs including 266 down-regulated and 449 up-regulated genes were identified. This included 18, 22 and 28 HICs, LMGs, and ISGs, respectively (Figure 2). A list of DEGs is provided in Supplementary File 1. Furthermore, of the total differentially expressed HICs, 7 were potassium channels: *KCNA6, KCNG3, KCNH4, KCNJ10, KCNJ12, KCNN3,* and *KCTD4.* Table 1 represents the list of identified dysregulated potassium channels.

**FIGURE 2.**
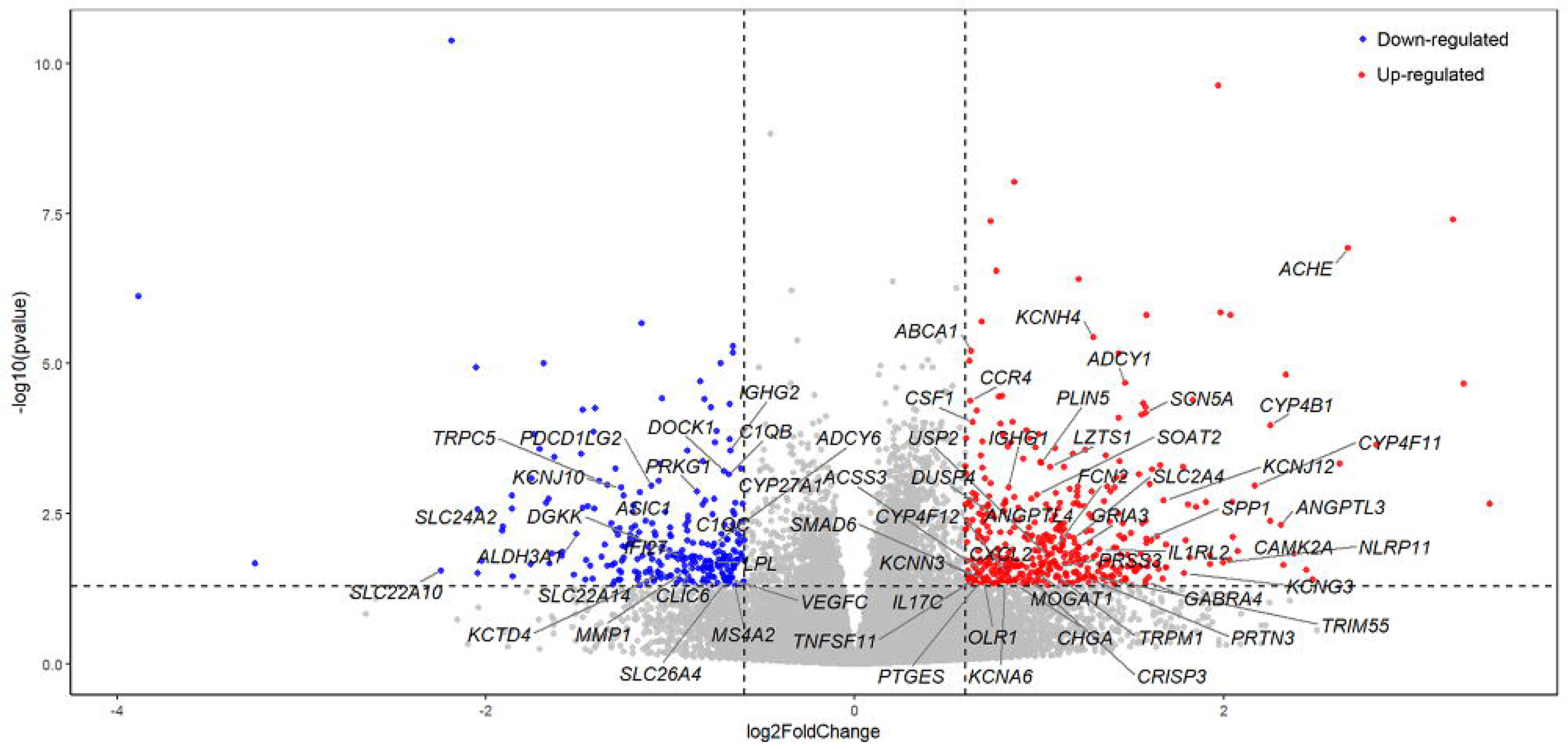
Volcano-plot displaying fold-change (Log2-FC) versus significance (-Log10 p-value). Red circles represent up-regulated genes (Log2-FC > 0.6) and blue circles represent down-regulated genes (Log2-FC < −0.6). Gray circles represent genes with |Log2-FC| < 0.6 or non-significant −Log10(p-value). Labels in the volcano plots specifically highlight differentially expressed human ion channels, lipid metabolism-, and immune signaling-related genes.

**Table 1:** Dysregulated potassium ion channels in LC patients. List of up- and down-regulated potassium channels in LC patients, with their corresponding description, log2Foldchange, significant p-value, and dysregulation status.

| Potassium channels | Description | log2FC | p-value | Dysregulation |
| --- | --- | --- | --- | --- |
| *KCNH4* | Potassium voltage-gated channel subfamily H member 4 | 1.29209 | 3.59E-06 | Up-regulated |
| *KCNJ12* | Potassium inwardly rectifying channel subfamily J member 12 | 1.67679 | 0.00191 | Up-regulated |
| *KCNG3* | Potassium voltage-gated channel modifier subfamily G member 3 | 1.78694 | 0.03089 | Up-regulated |
| *KCNN3* | Potassium calcium-activated channel subfamily N member 3 | 0.65056 | 0.03441 | Up-regulated |
| *KCNA6* | Potassium voltage-gated channel subfamily A member 6 | 0.81391 | 0.04806 | Up-regulated |
| *KCNJ10* | Potassium inwardly rectifying channel subfamily J member 10 | -1.26974 | 0.00116 | Down-regulated |
| *KCTD4* | Potassium channel tetramerization domain containing 4 | -1.05143 | 0.03132 | Down-regulated |

Functional enrichment analysis of DEGs led to the identification of 702 biological processes, 162 molecular functions, 77 cellular components, and 29 pathways in LC patients (Figure 3) (Supplementary File 2). Pathways including antigen processing and presentation, complement and coagulation cascades, biosynthesis of unsaturated fatty acids, N-glycan biosynthesis, 2-oxocarboxylic acid metabolism, phagosome, and neuroactive ligand‒receptor interaction were predicted to be suppressed in LC patients through their negative normalized enrichment score (NES). Similarly, pathways including type II diabetes mellitus, pancreatic secretion, and maturity onset diabetes of the young were found to be activated in LC patients through their positive NES.

**FIGURE 3.**
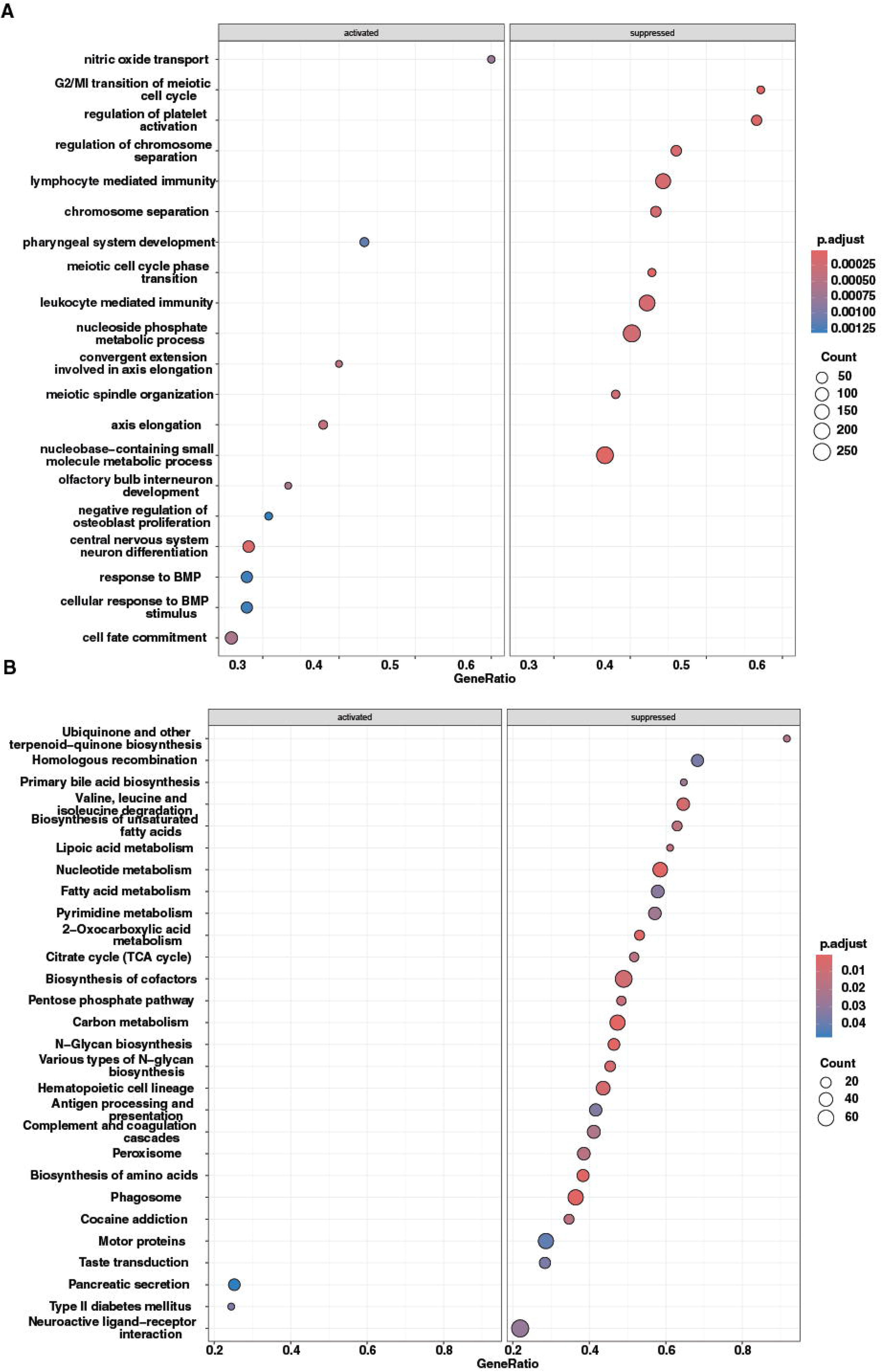
Functional enrichment of differentially expressed genes. A) Biological processes and B) pathways that were found to be activated and suppressed in long COVID patients.

### 3.2 Co-expression Network Analysis

Three significant co-expressed gene networks were identified using WGCNA, comprising the blue, brown, and turquoise modules, with 158, 98, and 79 DEGs, respectively. (Figure 4 and Supplementary File 3).

**FIGURE 4.**
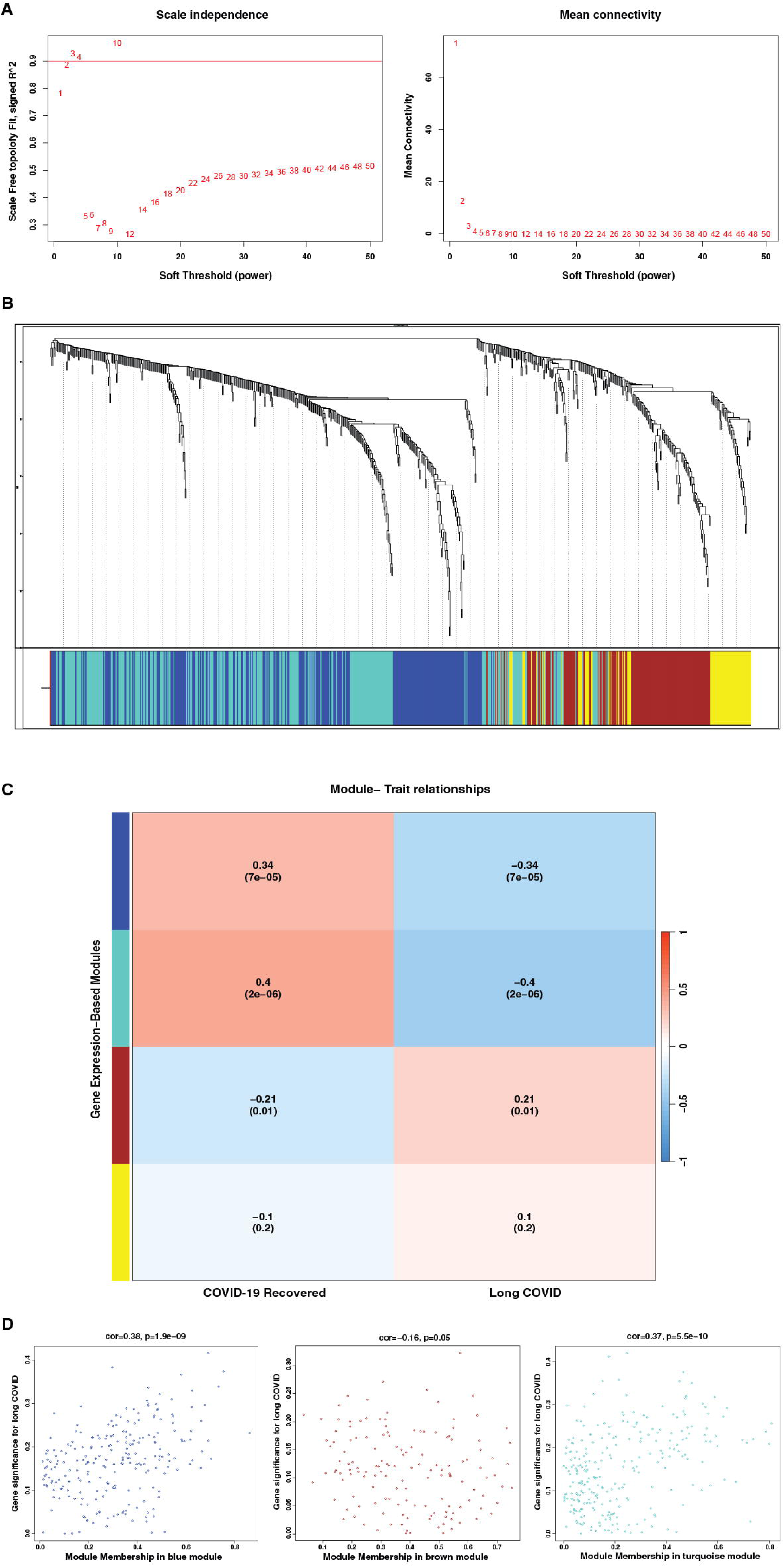
Weighted gene co-expression network analysis of dysregulated genes in long COVID (LC). (A) Scale independence plot depicting the relationship between soft threshold power (β=4) and scale-free topology fit. (B) Mean connectivity analysis illustrating how connectivity decreases with increasing soft threshold power. (C) Sample clustering dendrogram showing the relationship between COVID recovered and LC samples. (D) Module-trait relationship heatmap illustrating the correlation between identified modules and LC status, with red indicating a positive correlation and blue indicating a negative correlation. (E) Gene significance versus module membership plots for blue, brown and turquoise modules, demonstrating the relationship between gene significance for LC and intramodular connectivity.

The blue module comprised of 5 HICs - chloride intracellular channel 6 (*CLIC6*), potassium voltage-gated channel subfamily A member 6 (*KCNA6*), potassium inwardly rectifying channel subfamily J member 10 (*KCNJ10*), potassium calcium-activated channel subfamily N member 3 (*KCNN3*), and potassium channel tetramerization domain containing 4 (*KCTD4*); 7 LMGs - adenylate cyclase 6 (*ADCY6*), cytochrome P450 family 27 subfamily A member 1 (*CYP27A1*), cytochrome P450 family 4 subfamily B member 1 (*CYP4B1*), diacylglycerol kinase kappa (*DGKK*), lipoprotein lipase (*LPL*), matrix metallopeptidase 1 (*MMP1*), and protein kinase cGMP-dependent 1 (*PRKG1*); and 10 ISGs - complement C1q B chain (*C1QB*), complement C1q C chain (*C1QC*), dedicator of cytokinesis 1 (*DOCK1*), interferon alpha inducible protein 27 (*IFI27*), immunoglobulin heavy constant gamma 1 (G1m marker) (*IGHG1*), immunoglobulin heavy constant gamma 2 (G2m marker) (*IGHG2*), leucine zipper tumor suppressor 1 (*LZTS1*), membrane spanning 4-domains A2 (*MS4A2*), programmed cell death 1 ligand 2 (*PDCD1LG2*), and vascular endothelial growth factor C (*VEGFC*) (Figure 5A). The blue module was further found to be associated with several pathways, including the phospholipase D signaling pathway, complement and coagulation cascades, cell adhesion molecule (CAM) interaction, efferocytosis, and peroxisome proliferator-activated receptor signaling pathway.

**FIGURE 5:**
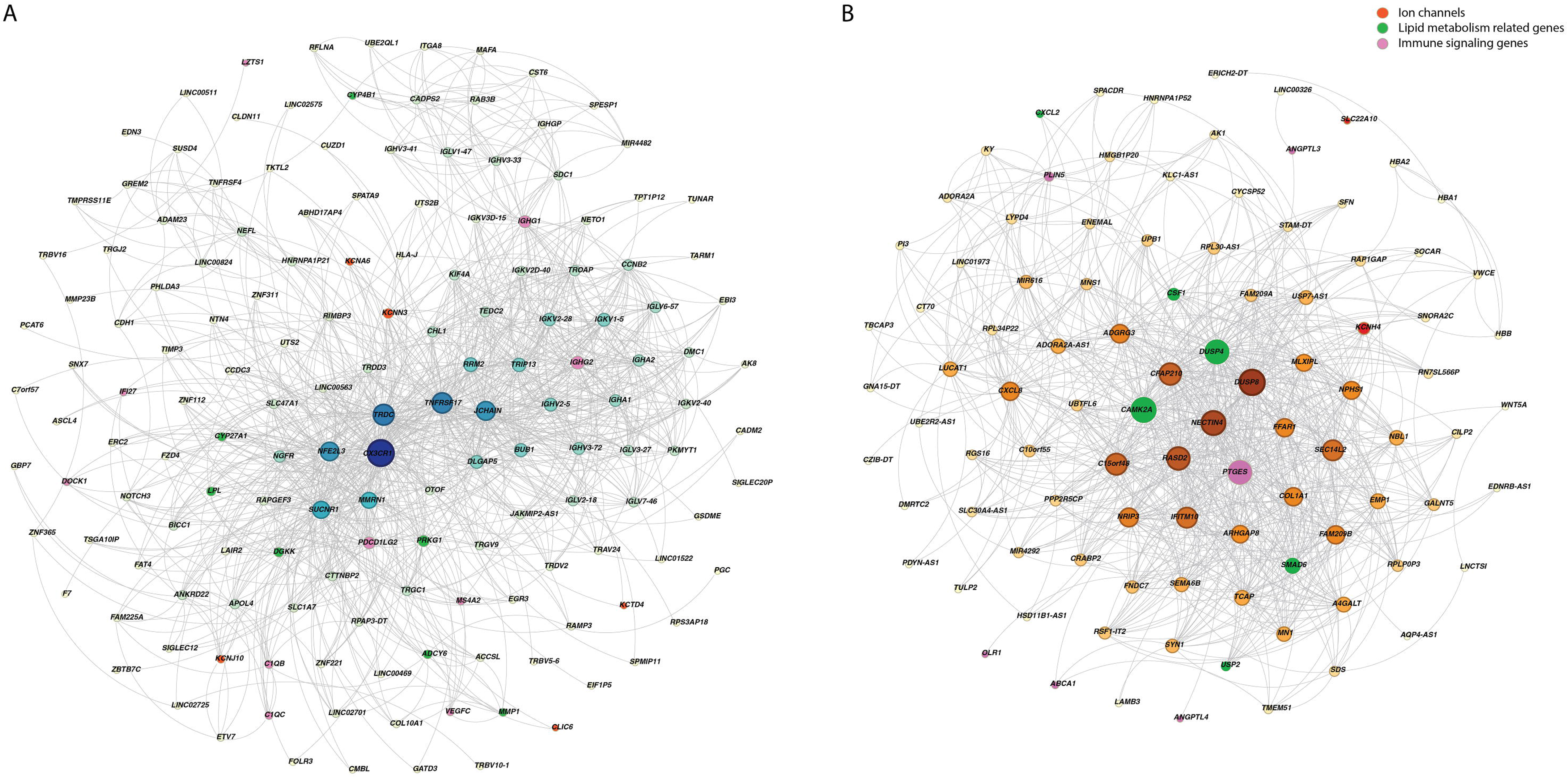
Co-expressed networks obtained through weighted gene co-expression network analysis: (A) Blue module visualization with ion channels highlighted in orange nodes, LMGs highlighted in green nodes, and ISGs highlighted in pink nodes. This module includes genes associated with the phospholipase D signaling pathway, complement and coagulation cascades, cell adhesion molecule (CAM) interaction, efferocytosis, and peroxisome proliferator-activated receptor signaling pathway. (B) Brown module visualization with channels highlighted in orange nodes, LMGs highlighted in green nodes, and ISGs highlighted in pink nodes. This module was further found to be associated with pathways including viral protein interactions with cytokine and cytokine receptor, lipid and atherosclerosis, and cholesterol metabolism.

The brown module contained 2 HICs that included potassium voltage-gated channel subfamily H member 4 (*KCNH4*) and solute carrier family 22 member 10 (*SLC22A10*), 6 LMGs - calcium/calmodulin dependent protein kinase II alpha (*CAMK2A*), colony stimulating factor 1 (*CSF1*), C-X-C motif chemokine ligand 2 (*CXCL2*), dual specificity phosphatase 4 (*DUSP4*), SMAD family member 6 (*SMAD6*), ubiquitin specific peptidase 2 (*USP2*), and 6 ISGs - ATP binding cassette subfamily A member 1 (*ABCA1*), angiopoietin like 3 (*ANGPTL3*), angiopoietin like 4 (*ANGPTL4*), oxidized low density lipoprotein receptor 1 (*OLR1*), perilipin 5 (*PLIN5*), and prostaglandin E synthase (*PTGES*) (Figure 5B). The brown module was further found to be associated with pathways including viral protein interactions with cytokine and cytokine receptor, lipid and atherosclerosis, cholesterol metabolism, among others.

The turquoise module contained 1 LMG – adenylate cyclase 1 (*ADCY1*) and 1 ISG – C-C motif chemokine receptor 4 (*CCR4*) and was found to be associated with gap junctions through pathway analysis.

### 3.3 Drug-target interactions

From the identified dysregulated HICs, 10 were found to interact with approved drugs (Supplementary File 4). Of these *KCNN3*, *KCNA6*, and *KCNJ10* were from the blue module, and *KCNH4* was from the brown module. *KCNN3* was observed to interact with dequalinium. *KCNJ10* interacted with mitiglinide, glipizide, tolazamide, and chlorpropamide. Additionally, both *KCNA6* and *KCNH4* were found to interact with amifampridine, guanidine hydrochloride, dalfampridine, and amifampridine phosphate. (Figure 6).

**FIGURE 6:**
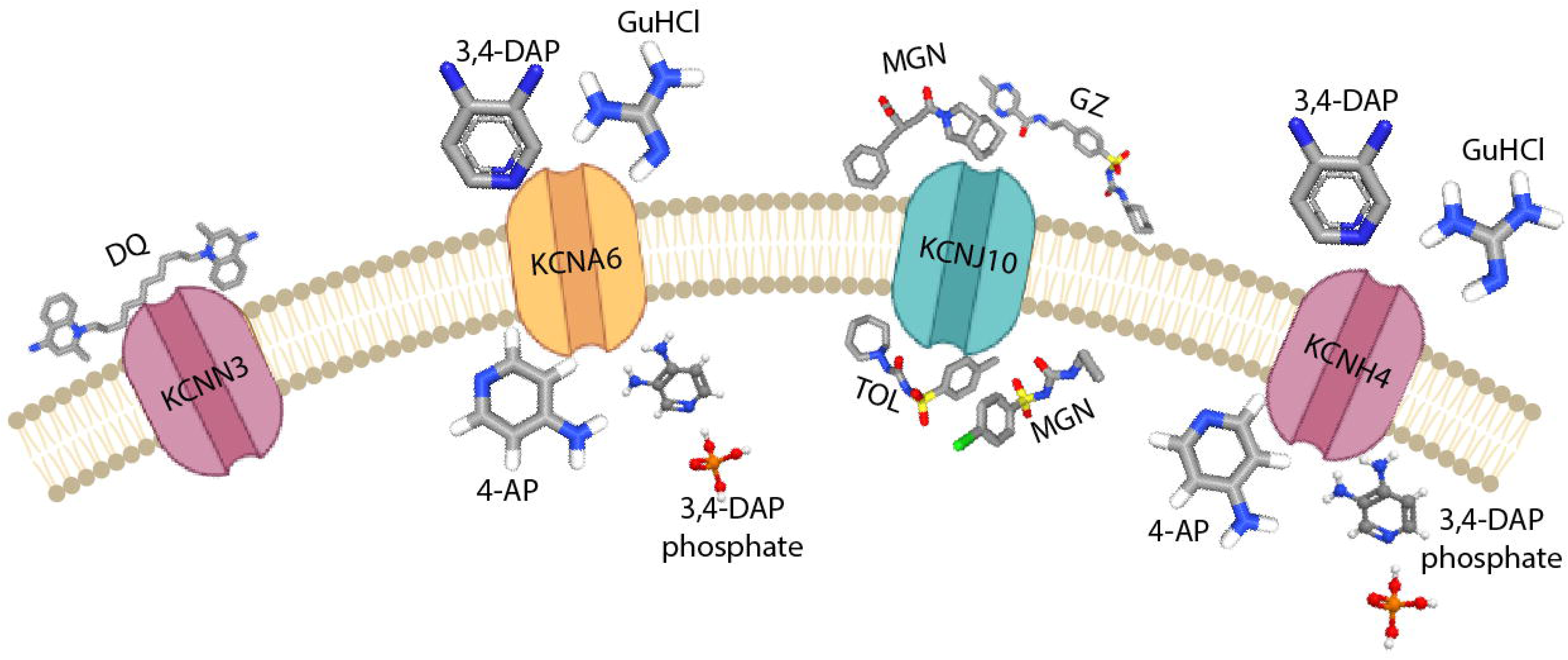
Interactions between potassium channels and drugs: (A) Potassium calcium-activated channel subfamily N member 3 (KCNN3) interacting with dequalinium. (B) Potassium voltage-gated channel subfamily A member 6 (KCNA6) interacting with amifampridine (3,4-DAP), guanidine hydrochloride (GuHCl), dalfampridine (4-AP), and amifampridine phosphate (3,4-DAP). (C) Potassium inwardly rectifying channel subfamily J member 10 (KCNJ10) is known to interact with mitiglinide (MGN), glipizide (GZ), tolazamide (TOL), and chlorpropamide. (D) Potassium voltage-gated channel subfamily H member 4 (KCNH4) is known to interact with Amifampridine (3,4-DAP), guanidine hydrochloride (GuHCl), dalfampridine (4-AP), and amifampridine phosphate (3,4-DAP).

Furthermore, eight statistically significant perturbagens (drugs/compounds) including etacrynic acid, lenalidomide, ciprofloxacin, naphazoline, exhibiting negative normalized connectivity scores were identified through the cMAP analysis, suggesting their potential to inversely modulate the LC-associated transcriptional signature (Table 2). Notably, dalfampridine, a potassium channel antagonist, was identified among the negatively connected perturbagens and targets several voltage-gated potassium channels, including *KCNA6*, *KCNG3*, and *KCNH4*, which were found to be dysregulated in LC patients. However, the fdr_q_nlog10 of dalfampridine was 0.0955 making it statistically non-significant.

**Table 2:** Perturbagens identified through Connectivity Map (cMAP) analysis of the LC gene expression signature. Perturbagens exhibiting negative normalized connectivity score indicate an inverse transcriptional relationship with the long COVID-associated gene expression signature and may represent potential candidates for transcriptome-based drug repurposing.

| Pert Identifier | Perturbagen (Drug/compound name) | Mode of Action (moa) | FDR_q_nlog10 | Normalized connectivity score |
| --- | --- | --- | --- | --- |
| BRD-K54256913 | MK-1775 | WEE1 kinase inhibitor | 15.3525 | -1.5824 |
| BRD-K63630713 | Etacrynic-acid | Sodium channel inhibitor | 15.6536 | -1.6201 |
| BRD-A04553218 | BRD-A04553218 | Histamine receptor antagonist | 15.6536 | -1.6459 |
| BRD-K22096725 | ALW-II-49-7 | Ephrin inhibitor | 15.3525 | -1.5966 |
| BRD-A17883755 | Lenalidomide | Carcinogen | 15.3525 | -1.5999 |
| BRD-K04804440 | Ciprofloxacin | Bacterial DNA Gyrase inhibitor | 15.6536 | -1.6043 |
| BRD-K77641333 | Naphazoline | Adrenergic receptor agonist | 15.6536 | -1.6788 |
| BRD-K93779381 | Ingenol-mebutate | PKC activator | 1.7758 | -1.5665 |
| BRD-K22482860 | Dalfampridine | Potassium channel antagonist | 0.0955 | -0.9713 |

### 3.4 Validation of Identified Differentially Expressed Genes in Independent Datasets

Differential gene expression analysis of the independent dataset led to the identification of 45 differentially expressed HICs, of which nine were potassium channels (*KCNJ6, KCNA6, KCND1, KCNH1, KCNH3, KCNJ15, KCNK7, KCNS1, KCNV2*). Among these 45 HICs - *KCNA6*, and gamma-aminobutyric acid type A receptor subunit alpha4 (*GABRA4*) were common with the identified differentially expressed HICs. Additionally, five LMGs (*ABCA1, CYP27A1, OLR1, PLIN5, ADCY1*) and one ISG (*CHGA*) were found to be dysregulated in both datasets. Furthermore, differential gene expression analysis of the independent longitudinal RNA-Seq dataset led to the identification of 47 differentially expressed HICs, of which 9 were potassium channels (*KCNC1, KCND3, KCNH2, KCNJ16, KCNJ6, KCNK10, KCNN1, KCNQ3, KCNT1*). Five HICs - *GABRA4*, solute carrier family 24 member 2 (*SLC24A2*), transient receptor potential canonical 5 (*TRPC5*), solute carrier family 26 member 4 (*SLC26A4*), and transient receptor potential melastatin 1 (*TRPM1*) were found to be dysregulated across both datasets.

## 4 Discussion

Given the latest definition of LC, the knowledge of mechanisms driving the condition and the multisystem nature of the symptoms experienced by the patients (54), (2), a meta-analysis of transcriptomic data from COVID-19-recovered and LC patients was performed to identify the underlying molecular alterations most likely responsible for these mechanisms.

Differential gene expression analysis led to the identification of DEGs in LC patients, including seven potassium ion channel genes, *KCNA6, KCNG3, KCNH4, KCNJ10, KCNJ12, KCNN3*, and *KCTD4,* among the differentially expressed HICs. Functional enrichment for the DEGs led to the identification of GO terms and pathways that may be altered in LC patients. Among the enriched GO terms, 171 biological processes, 47 molecular functions, and 21 cellular components included differentially expressed HICs, LMGs, and ISGs. Functional enrichment led to the identification of GO terms and pathways that are most likely activated or suppressed as a consequence of differential expression in LC patients. Pathways and processes related to diabetes and the nervous system were found to be activated, while several immune signaling, lipid metabolism, cell cycle, and other metabolic pathways were found to be suppressed in LC patients, as revealed by pathway enrichment analysis. Furthermore, significant modules of co-expressed genes that were associated with pathways linked with LC were identified. Notably, these module-associated pathways were consistent with those identified through initial pathway enrichment analysis, suggesting that genes within these modules may contribute to LC pathophysiology. Furthermore, of the seven potassium channels, five (*KCNN3*, *KCNH4*, *KCNA6*, *KCNJ10*, and *KCTD4*) were co-expressed in the identified modules and four of them (*KCNN3*, *KCNH4*, *KCNA6*, and *KCNJ10*) were found to interact with the approved drugs.

Dysregulated immune signaling and neurological pathways were found to be associated with LC in several transcriptomic studies. A transcriptomic data analysis of 48 LC patients reported 212 DEGs, including voltage-gated potassium channel subunit beta-1 (*KCNAB1*) (55). Additionally, whole-blood transcriptomic profiling identified sex-specific neuroinflammatory and immune dysregulation signatures in LC patients, including dysregulation of the potassium voltage-gated channel subfamily Q member 5 (*KCNQ5*), which was associated with immune dysregulation, cellular stress, and tissue repair (56). Notably, the identified HICs have also been implicated in independent transcriptomic and multi-omics investigations of post-viral and LC-associated pathophysiology.

To further assess the robustness of the findings, differential gene expression analysis using an independent RNA-Seq dataset was performed. This led to the identification of *KCNA6* and *GABRA4* common among the identified dysregulated HICs. Interestingly, both *KCNA6* and *GABRA4* were involved in regulating neurotransmitter release and their dysregulation was linked to neurodevelopmental disorders, epilepsy, and immune dysfunction (13), (57). Although the direction of dysregulation differed between the two datasets, its independent identification further strengthens the evidence supporting its association with LC-associated neurological manifestations. Nevertheless, the independent identification of *KCNA6* and *GABRA4* across transcriptomic datasets strengthens the evidence supporting its biological relevance in LC. Additionally, validation using a longitudinal RNA-Seq dataset comprising blood samples collected at 3–6 months and 12–15 months following SARS-CoV-2 infection further confirmed the differential expression of *SLC24A2*, *GABRA4*, *TRPC5*, *SLC26A4,* and *TRPM1*. Interestingly, the recurrent identification of *GABRA4*, a neurotransmitter regulator in the brain, across three independent studies, including the current meta-analysis, underscores its potential relevance in LC pathophysiology and highlights it as a promising neurologically associated LC marker

Prolonged exposure to TNF-α and IL-6 in human iPSC-derived neuron-astrocyte co-cultures were significantly dysregulated *KCNJ10*, leading to reduced Kir4.1-mediated potassium clearance and severe neural network dysfunction (58). These molecular changes result in impaired network activity and abnormal morphology, highlighting a mechanism for cognitive deficits in LC (58). Similarly, *KCNN3* has been identified in genomic and transcriptomic studies investigating cardiovascular and autonomic manifestations in LC and has been proposed as a candidate mediator of channelopathy-like late-onset complications following SARS-CoV-2 infection (59). Additionally, sodium voltage-gated channel alpha subunit 5 (*SCN5A*) has been reported in longitudinal multi-omics studies of LC, where its dysregulation has been associated with persistent tachycardia, palpitations, and cardiac conduction abnormalities that constitute common clinical manifestations of LC (60). Furthermore, acid-sensing ion channel subunit 1 (*ASIC1*) has been implicated in transcriptomic interactome studies examining SARS-CoV-2-mediated perturbations of ion channels and has been linked to chronic pain-related mechanisms, including mechanical hypersensitivity and nociplastic pain, phenotypes that overlap considerably with LC and myalgic encephalomyelitis/chronic fatigue syndrome (ME/CFS) (61).

*KCNN3*, belonging to the KCNN family of potassium channels, encodes a small conductance calcium-activated potassium channel (SK3). Functionally, SK3 mediates the voltage-independent transmembrane transfer of K^+^ across the cell membrane through its interaction with calmodulin, which binds the intracellular calcium, allowing its opening (10). Due to its higher expression in the brain and heart, dysregulation of *KCNN3* is reported to be associated with neurological disorders and cardiac arrhythmia, which are also among the manifestations observed in LC (62), (19), (63), (64). Dysregulated *KCNN3* has also been reported in previous bioinformatic studies, suggesting their involvement in LC-associated manifestations (65). SK3 is activated after membrane hyperpolarization, which then regulates neuronal excitability by contributing to the slow component of synaptic afterhyperpolarization (AHP). The identified upregulation of *KCNN3* may potentially contribute to delayed neuronal activation, a phenomenon observed in schizophrenia and atrial fibrillation, the latter of which has been reported to be associated with LC (66), (67). As a potential blocker of the SK3 channel, dequalinium prevents the slow reset period (AHP) followed by a nerve impulse, enhancing the excitability of a neuron (68), (66). Hence, the KCNN3-dequalinium interaction could provide a promising strategy in the management of neurological conditions observed in LC. Additionally, *KCNN3* was found to interact with 21 DEGs, of which 1 was LMG - protein kinase CGMP-dependent 1 (*PRKG1*) and 2 were ISGs - immunoglobulin heavy constant gamma 1 (*IGHG1*) and *PDCD1LG2*. Of them, *PDCD1LG2* has been reported as a major immune checkpoint during COVID-19 progression, while some have mentioned its elevation associated with T-cell exhaustion in LC cohorts (69), (70).

*KCNA6* is a voltage-gated potassium channel that is highly expressed in the central nervous system, with the highest expression in the frontal and occipital cortex (13). It regulates neuronal excitability by mediating potassium ion transport, which is crucial for repolarizing neurons after action potentials (71). Mutations in *KCNA6* disrupt channel deactivation, leading to persistent neuronal excitability and early-onset epilepsy, leading to behavioral, cognitive, and psychiatric issues (13), (72). Additionally, exogenous expression of *KCNA6* was found to significantly increase spike-mediated pseudoviral entry (21). Notably, disrupting KCNA6 channel activity has been shown to decrease SARS-CoV-2 viral entry, highlighting it as a putative SARS-CoV-2 host factor (21). The upregulation of *KCNA6* identified in this study, along with its interaction with channel blocker drugs pose them as a potential molecule in LC-associated neurological manifestations.

*KCNH4*, located in the neocortex and the striatum, is a brain-specific gene that is associated with the regulation of neurotransmitter release, heart rate, insulin secretion, neuronal excitability, epithelial electrolyte transport, smooth muscle contraction, and cell volume (73). Dysregulation of *KCNH4* is reported to be associated with sleep regulation, neurodegeneration, and several neurodevelopmental disorders. Several studies have reported sleep disturbances in patients with a history of acute COVID-19, suggesting that it is among the major neuropsychiatric symptoms experienced by patients (20), (74). Additionally, its co-expression with *PTGES*, an LMG and 2 ISGs, *CAMK2A* and *DUSP4,* was revealed through co-expression analysis. These molecules, especially *CAMK2A* and *DUSP4,* have been reported to be linked with SARS-CoV-2 viral entry and LC-associated brain fog, respectively (75), (76). *KCNH4* was identified to be upregulated. Furthermore, it was found to interact with potassium channel blockers including amifampridin.

Kir4.1, encoded by *KCNJ10*, is an inwardly rectifying potassium channel that maintains K^+^ homeostasis in glial cells in the central nervous system, inner ear cells, and renal epithelial cells (11). Specifically, it is involved in stabilizing the resting membrane potential of astrocytes, clears extracellular K^+^ to prevent neuronal over-excitation, and enables salt reabsorption in the kidney (12), (15). *KCNJ10* downregulation has been reported as a marker of astrocyte dysfunction in depression that may contribute to neuroinflammation-driven neuropsychiatric complications in LC (77). Reduced expression of *KCNJ10* contributes to loss of K^+^ buffering capacity and disruption of ionic equilibrium in retinal Müller glial cells (15), (78). Additionally, disruption in channel functionality has been reported to cause EAST syndrome, making it critical in neurological implications (79). Additionally, immune signaling involvement of *KCNJ10* through its co-expression with *PDCD1LG2* was also identified. The downregulation of *KCNJ10* in LC patients identified in the current analysis and its association with neurological imbalance reported in the literature highlight its potential relevance to neurological manifestations.

The identified potassium channel blockers are approved for the treatment of disease conditions including Lambert-Eaton myasthenic syndrome (LEMS) (amifampridine, amifampridine phosphate, and guanidine hydrochloride), multiple sclerosis (dalfampridine), bacterial infections (dequalinium), and type 2 diabetes mellitus (glipizide, mitiglinide, tolazamide, chlorpropamide) (80), (81), (82), (83), (84), (85). One of the potassium channel blockers - amifampridine, an FDA-approved drug for the treatment of LEMS-associated muscle weakness, blocks presynaptic potassium channels, thereby prolonging nerve action potentials and increasing calcium influx into nerve terminals (80), (86). This enhances acetylcholine release at the neuromuscular junction, resulting in improved neuromuscular transmission and muscle strength (80). Additionally, amifampridine has been reported as an experimental, off-label treatment to alleviate the severe muscular fatigue and post-exertional malaise associated with LC (87).

Similarly, dalfampridine, a voltage-gated potassium channel blocker, has been approved for the management of multiple sclerosis (85). Due to its lipid-soluble nature, dalfampridine readily crosses the blood-brain barrier (BBB) and may interact with several HICs, including KCNH4 and KCNA6, on the cytoplasmic side of neuronal membranes within the central nervous system (CNS) (85). In contrast, the limited lipid solubility of amifampridine restricts its BBB permeability, contributing to its efficacy and reduced CNS toxicity in the treatment of LEMS (86). Although BBB penetration may increase the risk of toxicity in certain contexts, it can also offer therapeutic advantages. For example, in the case of dequalinium, BBB penetration may facilitate drug delivery to intracranial tumors such as glioblastoma, where effective access to the brain is essential for therapeutic efficacy (88). This highlights the importance of considering BBB permeability when evaluating candidate compounds targeting the neurological manifestations of LC.

Interestingly, transcriptome-based perturbagen analysis using the cMAP identified dalfampridine as a negatively connected perturbagen, suggesting an inverse transcriptional relationship with the LC-associated gene expression signature. Dalfampridine is known to target several voltage-gated potassium channels, including KCNA6, KCNG3, and KCNH4. These ion channels were found to be dysregulated in patients with LC in this study. Identification of dalfampridine using DGIdb and cMAP analysis highlights potassium channel modulation as a potential pharmacological strategy for LC. However, dalfampridine did not satisfy the statistical significance threshold in the cMAP analysis (fdr_q_nlog10 = 0.0955) and exhibited a relatively modest normalized connectivity score (−0.9713). Therefore, its transcriptome-based association should be interpreted cautiously and considered hypothesis-generating for further experimental validation.

Several of the identified potassium channel blockers are known to exhibit pleiotropic pharmacological activities and may interact with molecular targets beyond the identified HICs, increasing the possibility of unintended pharmacological effects. Recent evidence suggests that amifampridine, in addition to blocking presynaptic potassium channels, can act as an agonist of L-type voltage-gated calcium channels at low millimolar concentrations, a mechanism that may further contribute to enhanced neurotransmitter release at the neuromuscular junction (89). This broad pharmacological activity has been associated with adverse effects including paresthesia, likely resulting from enhanced neurotransmission at autonomic and sensory nerve terminals. Furthermore, a generalized increase in acetylcholine release can stimulate the autonomic nervous system beyond the intended neuromuscular targets, leading to gastrointestinal adverse effects including abdominal pain, nausea, and diarrhea. Similar considerations regarding target specificity among other pharmacological effects may apply to the other identified drugs.

## 5 Limitations

The bulk RNA-Seq transcriptomic data obtained from blood-derived samples, that capture gene expression patterns but do not directly verify protein-level changes or functional modifications in HICs, LMGs, and ISGs were analyzed. However, it provides the first line of evidence for several future studies to establish a definitive link between translation and LC. Furthermore, the integrated datasets included different peripheral blood-derived sample types, including whole blood and peripheral blood mononuclear cells, which differ in their cellular composition and may contribute to biological and technical variability across studies. Although standardized preprocessing and batch-effect correction approaches were employed to minimize inter-study variability, residual heterogeneity arising from differences in sample types and patient characteristics cannot be completely excluded. Moreover, the heterogeneity of LC progression across individuals and differences in the underlying disease mechanisms may constrain the generalizability of these findings. Although independent transcriptomic validation was performed in an additional post-COVID-19 cohort, the availability of publicly accessible longitudinal RNA-Seq datasets with serial sampling remains limited, preventing assessment of the temporal stability and progression of HIC dysregulation in LC. Future studies incorporating larger, longitudinal patient cohorts, single-cell transcriptomic approaches, with experimental validation would not only strengthen these findings but also provide more insights into the complex pathophysiology underlying LC.

Secondly, identified drugs interacting with altered potassium channels should be considered preliminary and hypothesis-generating. Although these findings provide a basis for exploring potassium channel-targeted therapeutic strategies in LC, the efficacy of the identified drugs remains to be established in this disease context. Additional experimental and clinical studies are required to validate these drug-target interactions and their downstream biological consequences before these compounds can be considered viable candidates for the better management of patients with LC.

## 6 Conclusion

The significantly co-expressed networks, and altered pathways associated with neurological, renal, and cardiovascular processes were identified through computational analysis and extensive data mining approach. Notably, the dysregulation and co-expression of HICs, particularly potassium channels, highlight their potential in LC management. Collectively, these results suggest a critical subset of molecular alterations underlying broad multisystem dysregulation in LC and provide a molecular framework for understanding its pathophysiology, thereby facilitating the identification of potassium channels as potential targets. However, further experimental validation in larger cohorts is necessary to confirm these observations and translate them into effective diagnostic and treatment strategies.

## CRediT Authorship Contribution Statement

John P. George: Writing – review & editing, Writing – original draft, Visualization, Methodology, Investigation, Formal analysis, Data curation. Kiran Bharat Gaikwad: Writing – review & editing, Visualization, Investigation. Jyoti Sharma: Writing – review & editing, Supervision, Methodology, Investigation, Funding acquisition, Conceptualization.

## Supporting information

Supplementary Filles 1,2,3,4,5,6

## Acknowledgements

We would like to thank the authors of the manuscripts for making the datasets used in this study publicly available. J.S. would like to thank Council of Scientific and Industrial Research (CSIR), Government of India [37WS(0114)/2023-24/EMR-II/ASPIRE] and Indian Council of Medical Research (ICMR), Government of India [FIW-2025-01-00001091] for the research support. J.P.G. is supported by the CSIR [37WS(0114)/2023-24/EMR-II/ASPIRE], and K.B.G. is supported by the ICMR [FIW-2025-01-00001091]. J.S. was a recipient of the Bio-CARe Women Scientist award from the Department of Biotechnology (DBT), Government of India.

## Data Availability Statement

The datasets used in this study are available in NCBI’s Gene Expression Omnibus database (https://www.ncbi.nlm.nih.gov/geo/) and BioProject (https://www.ncbi.nlm.nih.gov/bioproject/). These datasets were derived using the following accessions - GSE251849, GSE226260, and PRJNA1184005. R script used for differential gene expression is available at https://github.com/js-iob/John_CovRec_longCovid/blob/main/CocRec_vs_LC_processing_and_analysis_032626.R. R script used for functional enrichment is available in Github repository at https://github.com/js-iob/John_CovRec_longCovid/blob/main/Functional_enrichment_analysis_041026.R. R script used for WGCNA is available in Github repository through this link: https://github.com/js-iob/John_CovRec_longCovid/blob/main/WGCNA_to_network.R.

## Supplementary Data

***Supplementary File 1***. Differentially expressed genes identified in LC patients compared with COVID-19-recovered individuals

***Supplementary File 2***. Functional enrichment analysis results obtained using ClusterProfiler and Enrichr

***Supplementary File 3***. Node and edge information for the three significant WGCNA modules (Blue, Brown, and Turquoise)

***Supplementary File 4***. Drug interacting with the identified differentially expressed human ion channels retrieved from the Drug-Gene Interaction Database

***Supplementary File 5.*** Connectivity Map (cMAP) query results for the gene expression signature identified in patients with long COVID as compared with COVID-19-recovered

***Supplementary File 6.*** List of R packages and tools/utilities with version information used in this study

